# A comprehensive collection of transcriptome data to support life cycle analysis of the poplar rust fungus *Melampsora larici-populina*

**DOI:** 10.1101/2021.07.25.453710

**Authors:** Pamela Guerillot, Emmanuelle Morin, Annegret Kohler, Clémentine Louet, Sébastien Duplessis

## Abstract

*Melampsora larici-populina* is an obligate biotrophic plant pathogen responsible for the poplar rust disease. This fungus belongs to the taxonomical order Pucciniales and exhibits a complex heteroecious and macrocyclic life cycle, i.e. it has the capacity to infect two unrelated host plants, larch and poplar, and to form five distinct spore types through the year. The *M. larici-populina* genome has been sequenced and annotated in 2011 and since, different transcriptomic analyses were conducted at different stages with oligoarrays and later on with RNA-Seq covering most of its life cycle. Here, we collected published transcriptome data for the poplar rust fungus, transposed them for the version 2 of the genome now available at the Joint Genome Institute Mycocosm website and performed normalization of different oligoarray datasets on one hand, and of RNA-Seq on the other hand. We report a comprehensive normalized dataset for this fungus, made available for the community, which allows life cycle transcriptomics analysis.

## 1. Research background

Rust fungi belong to the taxonomical order Pucciniales that consists of more than 7000 species (Aime and McTaggart, 2021). These obligate biotrophic fungi infect a large number of host plants in which they cause important damages. They are a threat to food security and their repeated epidemics are under surveillance worldwide (Figueroa et al. 2017, Lorrain et al. 2019). Since their trophic status impedes easy manipulation under laboratory conditions -i.e. they must be multiplicated in living host plants and there are no routine genetic manipulations-genomics and transcriptomics have been approaches of choice which have allowed great progress in the past ten years in the understanding of their biology (Aime et al. 2017; Bakkeren and Szabo, 2020). Rust fungi exhibit the most complex life cycle among fungi with (i) the capacity to alternate onto two different taxonomically unrelated host plants (called heteroecism in such rust fungi), (ii) the production of up to five different spore types through the year (called macrocyclic rust fungi) and (iii) the presence of a dormancy period during the winter season for rust fungi found in temperate, montane or boreal conditions (Duplessis et al. 2021). Despite such an intriguing biological cycle marked by many different developmental and environmental controlled stages, most of the transcriptomics studies conducted in rust fungi were limited to a few conditions (Duplessis et al. 2012; Duplessis et al. 2014). Indeed, the majority of the transcriptome studies were focused on the host in which epidemics and major symptoms are recorded with time-course infection studies, isolation of infection structures or spore germination and host colonization (Lorrain et al. 2019). Only recently, a larger number of biological stages were considered for transcriptomics in a handful of rust fungi (for a recent review, see Duplessis et al. 2021).

*M. larici-populina* is responsible for the poplar rust disease and causes important loss in poplar plantations worldwide (Steenackers et al. 1996). It was a pioneer target for rust genomics (Duplessis et al. 2011a). Several transcriptome analyses were performed in the last decade aiming at studying genetic programs expressed at specific stages of the life cycle, and particularly in the different spores that are produced on the two host plants, larch and poplar (Duplessis et al. 2021).

As more genomes are made available within the fungal order Pucciniales, more transcriptome studies will be carried and comparative transcriptomics through cross-comparison of datasets between species will foster a better understanding of the biology of these fungi. Our goal here is to provide a comprehensive consolidated dataset of life cycle expression profiling for the poplar rust *M. larici-populina* that is up-to-date with the most recent version of the genome (version 2) available at the JGI Mycocosm, that can be easily explored and compared with other rust. Moreover, rust fungal genomes exhibit specific genomic features such as a large genome size, a high content in transposable elements and expanded gene catalogs (Aime et al. 2017). Altogether, these features render genome sequencing and assembly tedious and time-consuming, even with long molecule sequencing technologies. Thus, transcriptomics of rust species without a reference genome is still a favorite approach and more analysis may be expected in the coming years. Comparison to the poplar rust genome annotation and to the life cycle transcriptomics dataset reported here will allow researchers in the rust community to rapidly compare their data and strengthen their hypotheses.

## 2. Methodology

### Preliminary note

*Although all transcriptomics data were produced in our laboratory, we decided to realize data collection and comparison as if we were an external user, collecting published expression datasets from the NCBI transcriptome repository Gene Expression Omnibus. Additional datasets were deposited in the FigShare repository (doi: 10.6084/m9.figshare.15048822). In case of discontinuity of service, the datasets are available from the corresponding author.*

### Original *M. larici-populina* transcriptomic datasets

Transcriptomics data established with the first version of the *M. larici-populina* reference genome isolate 98AG31 were retrieved from the NCBI Gene Expression Omnibus portal (https://www.ncbi.nlm.nih.gov/geo/) using source data identifiers GSE21624, GSE49099, and GSE106863 described in the related publications: dormant and germinated urediniospores and time course infection of poplar leaves from 24 to 168 hours post-inoculation (hpi) by urediniospores (Nimblegen oligoarrays; Duplessis et al., 2011b); telia collected in autumn before winter dormancy (Nimblegen oligoarrays; Hacquard et al., 2013); basidia produced from telia right after exiting dormancy, and pycnia and aecia produced on larch needles after infection with basidiospores (Illumina HiSeq RNA-Seq; Lorrain et al., 2018). All RNA sources correspond to the genome reference isolate 98AG31, except for telia, which were isolated from natural samples (Hacquard et al., 2013).

### Processing of oligoarray expression datasets

In the original publications and GEO series, expression profiles were established based on the gene annotation of *M. larici-populina* genome version 1 (16,499 genes). The current genome annotation (version 2) contains 19,550 genes, of which only a portion overlaps, making direct comparison limited. Genome annotations are available from the JGI Mycocosm website (https://mycocosm.jgi.doe.gov; Grigoriev et al. 2014). The initial expression data were re-analyzed in an attempt to transpose available expression data to the version 2 of the poplar rust genome annotation. First, the oligonucleotide sequences from the custom-commercial Roche-NimbleGen arrays (GEO platform GPL10350) were compared to the transcripts annotated in the version 2 of the genome using the blastn tool and were assigned to a given transcript only if the sequence showed a strict identity of 100% with a given oligonucleotide. Out of 72,038 oligoprobes, 50,676 matched to a transcript from the version 2 (with 1 to 4 probes per transcript). These 50,676 oligoprobes were associated with 13,804 transcripts from the version 2 of the *M. larici-populina* genome. These oligoprobes were selected along with 1,064 random probes used to determine background levels and are presented in the Dataset 1. Then, the raw fluorescence values assigned to the oligonucleotides in the GSE21487 and GSE49099 series were retrieved from the GEO website and all these data were analyzed with the ARRAYSTAR software (DNASTAR, Inc. Madison, WI, USA) following the procedure described in the original publications. ARRAYSTAR generates a normalization of the expression level of each transcript across oligoprobes and across biological replicates of the different biological stages considered. The major difference here with the original publications, is the normalization of all samples from Duplessis et al. (2011b) and Hacquard et al. (2013) altogether. The normalized expression values of the 13,804 transcripts are presented in the Dataset 2.

### Processing of RNA-Seq expression datasets

Illumina RNA-Seq raw data from GSE106863 (Lorrain et al. 2018) were re-analyzed using the *M. larici-populina* genome version 2 following a similar procedure than in the original publication and the updated RNA-Seq analysis is now available at GEO as series GSE180724. Briefly, paired end reads from *M. larici-populina* basidia, spermogonia (also called pycnia) and aecia were mapped to transcripts from the poplar rust genome version 2 using the CLC Genomics Workbench 12.0 software (QIAGEN Digital Insights). Based on the CLC quality control, five nucleotides were removed at the 5′ and 3′ ends of the reads and the trimmed sequences were mapped onto *M. larici-populina* transcripts (https://mycocosm.jgi.doe.gov/Mellp2_3/; Mellp2_3_GeneCatalog_transcripts_20151130.nt.fasta) using the CLC RNA-Seq Analysis procedure considering 90% identity over 90% of the sequence length (10 hits maximum per read). The unique reads values were retrieved for each transcript and data analysis and normalization were performed with DESeq2 (v1.32.0; Love et al. 2014) as described in Lorrain et al. 2018. Unique reads obtained after the CLC RNA-Seq analysis are available in the series GSE180724 and all normalized read counts obtained from DESeq2 are available in the Dataset 3. The R program version 4.1.0 (R core team, 2020) was used for presenting distribution of expression data in boxplots, through the packages tidyverse v1.3.1 (Wickham et al. 2019) and ggplot2 v3.3.5 (Wickham, 2009).

## 3. M. larici-populina life cycle transcriptomics dataset description

Expression values obtained with NimbleGen oligoarrays and RNA-Seq for the 13,804 transcripts annotated in the *M. larici-populina* genome version 2 were combined into a single Dataset 4 in order to view expression levels at the following life cycle stages: dormant and germinated urediniospores, time course infection of poplar leaf tissues by urediniospores at 24, 48, 96 and 168 hours post-inoculation (hpi), early telia in autumn, basidia formed in spring, and spermogonia (pycnia) and aecia produced after 7 and 14 days on larch needles. The box-plots in Figure 1 show the distribution of expression data at the different stages. Attempts were made to cross normalize the oligoarrays and RNA-Seq data (e.g. quantile normalization) to help comparison, however low expression values close to (or below) background systematically artificially rose at too high levels for accurate use of the data (not shown). The best option remained to place the life stages expression levels next to each other with the knowledge of the differences and limits in the methods used for data generation. The Table 1 shows the levels of the quartiles from the box-plots displayed in Figure 1, which allows to determine whether the expression levels in the dataset should be considered below background, very low (below quartile 1), medium low or medium high (between the median and adjacent quartiles 1 and 3, respectively), or high (between quartile 3 and maximum value). Since cross-normalization was not satisfying, while exploring Dataset 4, users must always have in mind the intrinsic nature of the data and the distribution of expression values as depicted in Figure 1.

**Table 1.** Distribution of normalized expression levels of the *Melampsora larici-populina* life cycle transcriptomics dataset.

|  | Oligorrays |  |  |  |  |  |  | RNA-Seq |  |  |
| --- | --- | --- | --- | --- | --- | --- | --- | --- | --- | --- |
| Life stages <sup>a</sup> | URE | UREg | T24 | T48 | T96 | T168 | TEL | BSD | SPG | AEC |
| Background <sup>b</sup> | 18 | 18 | 44 | 30 | 19 | 16 | 16 | 9 | 6 | 20 |
| Min. | 15 | 15 | 17 | 14 | 15 | 14 | 14 | 1 | 1 | 1 |
| Quartile q1 | 38 | 36 | 165 | 93 | 49 | 36 | 36 | 2 | 3 | 2 |
| Median | 309 | 350 | 526 | 425 | 295 | 366 | 360 | 196 | 138 | 158 |
| Quartile q3 | 3,377 | 3,376 | 1,615 | 1,954 | 2,989 | 3,177 | 3,275 | 2,425 | 904 | 1,130 |
| Max. | 42,536 | 42,106 | 62,884 | 57,820 | 51,760 | 47,344 | 48,967 | 1,175,964 | 653,273 | 541,268 |

<sup>a</sup> URE, dormant urediniospores; UREg, germinated urediniospores; T24, T48, T96 and T168, hours post-inoculation during time course infection of poplar leaves; TEL, telia; BSD, basidia; SPG, spermogonia at 1 week post-inoculation on larch needles; AEC, aecia at 2 weeks post inoculation on larch needles.
<sup>b</sup> background levels were established from 1063 random oligonucleotides on the Nimblegen custom oligoarrays and background count values for RNAseq were considered at an arbitrary level of 10 unique reads.

**Figure 1.**
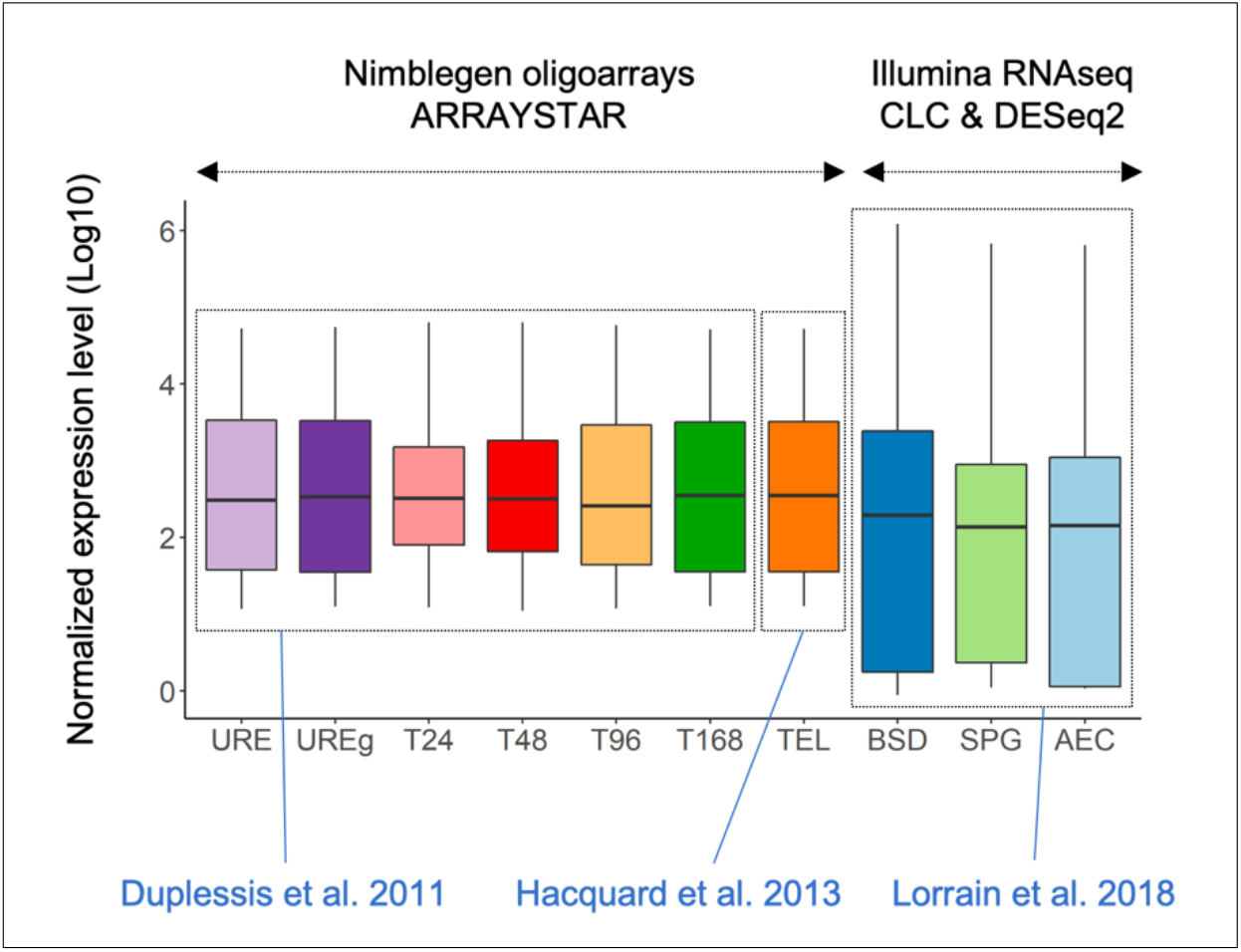
Distribution of normalized expression levels at various stages of the *M. larici-populina* life cycle. Boxplots showing the distribution of average normalized expression values (fluorescence levels or sequencing counts). The top and bottom of boxes correspond to the 25 and 75 % quartiles, respectively. The middle line represents the median. Bottom and top lines refer to the min and max values respectively. The source of the data, i.e. publication, transcriptome approach and softwares, are indicated below and above the box-plots, respectively.

### Datasets

Dataset 1 to 4 are available in the FigShare repository under doi: 10.6084/m9.figshare.15048822.

#### Dataset 1

Match between oligonucleotides of the GEO platform GPL10350 and genes from the *M. larici-populina* genome version 2.

#### Dataset 2

Normalized expression values of 13,804 transcripts in the *M. larici-populina* genome version 2 detected at life cycle stages initially reported in Duplessis et al. (2011b) and Hacquard et al. (2013) from oligoarray-based transcriptomics.

#### Dataset 3

Normalized expression values of 19,550 genes from *Melampsora larici-populina* genome version 2 determined from life cycle stages reported in Lorrain et al. (2018) from RNA-Seq transcriptomics.

#### Dataset 4

*Melampsora larici-populina* life cycle transcriptomics dataset. Expression values of 13,804 genes in the *M. larici-populina* genome version 2 determined with oligoarrays and RNA-Seq experiments reported in Duplessis et al. (2011a), Hacquard et al. (2013) and Lorrain et al. (2018).

## Acknowledgements

AK, CL, EM and SD are supported by the French “Investissement d’Avenir” program ANR-11-LABX-0002-01, Lab of Excellence ARBRE. CL is supported by a PhD fellowship from the Region Lorraine and the French National Research Agency (ANR-18-CE32-0001, Clonix2D).

## Notes

https://doi.org/10.6084/m9.figshare.15048822.v1

https://www.ncbi.nlm.nih.gov/geo/query/acc.cgi?acc=GSE180724

### Competing Interest Statement

The authors have declared no competing interest.

## References

Aime MC, MacTaggart AR, Mondo SJ, Duplessis S. 2017. Phylogenetics and phylogenomics of rust fungi. Adv. Fungal Genet. 100: 267–307.

Aime MC, McTaggart AR. 2021. A higher-rank classification for rust fungi, with notes on genera. Fungal Syst. Evol. 7: 21–47.

Bakkeren G, Szabo LJ. 2020. Progress on molecular genetics and manipulation of rust fungi. Phytopathology 110: 532–543

Duplessis S, Cuomo CA, Lin YC, Aerts A, Tisserant E, Veneault-Fourrey C, Joly DL, Hacquard S, Amselem J, Cantarel B et al. 2011a. Obligate biotrophy features unravelled by the genomic analysis of rust fungi. Proc. Natl. Acad. Sci. USA 108: 9166–9171.

Duplessis S, Hacquard S, Delaruelle C, Tisserant E, Frey P, Martin F, Kohler A. 2011b. *Melampsora larici-populina* transcript profiling during germination and timecourse infection of poplar leaves reveals dynamic expression patterns associated with virulence and biotrophy. Mol. Plant–Microbe Interact. 24: 808–818.

Duplessis S, Joly DJ, Dodds PN. 2012. Rust effectors. In: Martin F, Kamoun S, eds. Effectors in plant-microbes interactions. Oxford, UK: Wiley-Blackwell, 155–193.

Duplessis S, Bakkeren G, Hamelin R. 2014. Advancing knowledge on biology of rust fungi through genomics. Adv. Bot. Res. 70: 173–209.

Duplessis S, Lorrain C, Petre B, Figueroa M, Dodds P, Aime MC. 2021. Host Adaptation and virulence in heteroecious rust fungi. Annu. Rev. Phytopathol. 59: 17.

Figueroa M, Hammond-Kosack KE, Solomon PS. 2017. A review of wheat diseases: a field perspective. Mol. Plant Pathol. 19: 1523–1536.

Grigoriev IV, Nikitin R, Haridas S, Kuo A, Ohm R, Otillar R, Riley R, Salamov A, Zhao X, Korzeniewski F, et al. 2014. MycoCosm portal: gearing up for 1000 fungal genomes. Nucleic Acids Res. 42: D699–D704.

Hacquard S, Delaruelle C, Frey P, Tisserant E, Kohler A, Duplessis S. 2013. Transcriptome analysis of poplar rust telia reveals overwintering adaptation and tightly coordinated karyogamy and meiosis processes. Frontiers Plant Sci. 4: 456.

Lorrain C, Marchal C, Hacquard S, Delaruelle C, Petrowski J, Petre B, Hecker A, Frey P, Duplessis S. 2018. The rust fungus *Melampsora larici-populina* expresses a conserved genetic program and distinct sets of secreted protein genes during infection of its two host plants, larch and poplar. Mol. Plant–Microbe Interact. 31: 695–706.

Lorrain C, Gonçalves Dos Santos KC, Germain H, Hecker A, Duplessis S. 2019. Advances in understanding obligate biotrophy in rust fungi. New Phytol. 222: 1190–1206.

Love MI, Huber W, Anders S. 2014. Moderated estimation of fold change and dispersion for RNA-seq data with DESeq2. Genome Biol. 15: 550.

R Core Team (2020).R: A language and environment for statistical computing. R Foundation for Statistical Computing, Vienna, Austria. URL: http://www.r-project.org/index.html.

Steenackers J, Steenackers M, Steenackers V, Stevens M. 1996. Poplar diseases, consequences on growth and wood quality. Biomass Bioenerg. 10: 267–274.

Wickham H, Averick M, Bryan J, Chang W, D’Agostino McGowan L, François R, Grolemund G, Hayes A, Henry L, et al. 2019. Welcome to the tidyverse. J. Open Source Software 4: 1686.

Wickham H. 2009. ggplot2: Elegant Graphics for Data Analysis. Springer-Verlag, New York.

